# Specificity of RNAi, LNA and CRISPRi as loss-of-function methods in transcriptional analysis

**DOI:** 10.1101/234930

**Authors:** Lovorka Stojic, Aaron Lun, Jasmin Mangei, Patrice Mascalchi, Valentina Quarantotti, Alexis R Barr, Chris Bakal, John C Marioni, Fanni Gergely, Duncan T Odom

**Author notes:** Equal contributions (Lovorka Stojic, Aaron Lun). Present Address: Patrice Mascalchi: Bordeaux Imaging Center, UMS 3420 CNRS, US4 INSERM, University of Bordeaux, 33000 Bordeaux, France. Jasmin Mangei: German Cancer Research Center (DKFZ), Division: Molecular Genetics, Im Neuenheimer Feld 580, D-69120 Heidelberg.

## Abstract

Loss-of-function (LOF) methods, such as RNA interference (RNAi), antisense oligonucleotides or CRISPR-based genome editing, provide unparalleled power for studying the biological function of genes of interest. When coupled with transcriptomic analyses, LOF methods allow researchers to dissect networks of transcriptional regulation. However, a major concern is nonspecific targeting, which involves depletion of transcripts other than those intended. The off-target effects of each of these common LOF methods have yet to be compared at the whole-transcriptome level. Here, we systematically and experimentally compared non-specific activity of RNAi, antisense oligonucleotides and CRISPR interference (CRISPRi). All three methods yielded non-negligible offtarget effects in gene expression, with CRISPRi exhibiting clonal variation in the transcriptional profile. As an illustrative example, we evaluated the performance of each method for deciphering the role of a long noncoding RNA (lncRNA) with unknown function. Although all LOF methods reduced expression of the candidate lncRNA, each method yielded different sets of differentially expressed genes upon knockdown as well as a different cellular phenotype. Therefore, to definitively confirm the functional role of a transcriptional regulator, we recommend the simultaneous use of at least two different LOF methods and the inclusion of multiple, specifically designed negative controls.

## INTRODUCTION

The ability to specifically reduce the expression of a particular coding or noncoding gene is fundamental for establishing its loss-of-function phenotype in cells and organisms, and is frequently the only way to infer its function. Three popular strategies for doing this are RNA interference (RNAi), locked nucleic acid (LNA) hybridization and the CRISPR/Cas9 system (1). RNAi uses small interfering RNA oligonucleotides to deplete target transcripts by triggering their degradation (2). Efficient depletion of genes can also be achieved with antisense oligonucleotides (3); the most widely used antisense approach is locked nucleic acids (4), which can trigger RNase-H-mediated degradation of the target in the nucleus (5).

Most recently, the CRISPR/Cas9 system has been adapted to inhibit the expression of single genetic locus. Deactivated Cas9 (dCas9) fused to a Krüppel-associated box (KRAB) repression domain can be directed to a specific genomic locus to prevent transcription, an approach known as CRISPR interference (CRISPRi) (6-9). This allows repression of a targeted locus without editing the genome, thus avoiding unintentional deletion of active regulatory elements (10). CRISPRi can either be applied in homogenous populations derived from single cells expressing dCas9-KRAB or in poly- or non-clonal populations (11,12). This approach can successfully deplete both protein-coding genes (13) and noncoding RNAs such as long noncoding RNAs (lncRNAs) (8,14-16) and miRNAs (17). The apparent high specificity of CRISPRi has made it a preferred alternative to conventional gene silencing methods, particularly for lncRNAs, which are frequently located in complex genomic landscapes (10) (14).

Although all of these strategies have been successfully applied against a wide variety of targets in a range of biological systems, a critical consideration is whether the expression levels of non-target genes are inadvertently perturbed. These off-target effects arise from non-specific activity of the knockdown technology when exposed to the endogenous pool of total RNAs. For example, RNAi and antisense oligonucleotides with sufficient complementarity to non-target transcripts may cause unintended repression of those genes (18-21). Widespread genome binding and modest offtarget effects have also been reported for the dCas9-KRAB system (22,23).

In this study, we comprehensively quantify the off-target effects associated with each loss-of-function (LOF) strategy, with a particular focus on the transcriptome. Exploiting HeLa cell line as a powerful and easily modifiable model, we performed RNA sequencing (RNA-seq) with a range of negative controls for each method and used differential expression analyses to identify the off-target effects. RNAi and LNA approaches were notably dependent on the precise sequences of in the siRNA/antisense oligonucleotides used in the negative controls. The impact of introducing dCas9-KRAB to generate a polyclonal population of HeLa cells caused few transcriptional perturbations; in contrast, the process of single cell cloning from this population resulted in unique transcriptional signatures. To illustrate the impact that differences between methods can have on understanding gene function, we describe how the different LOF methods used to deplete a nuclear lncRNA can lead to different biological conclusions.

## MATERIAL AND METHODS

### Cell culture

HeLa and HEK293T cells were maintained in Dulbecco’s modified Eagle’s medium (Sigma Aldrich, D6429) supplemented with 10% Fetal bovine serum (FBS, Thermo Fisher Scientific). Both cell lines were obtained from American Type Culture Collection (ATCC) and were cultured at 37°C with 5% CO_2_. HeLa Kyoto (EGFP-α-tubulin/ H2B-mCherry) cells were obtained from Jan Ellenberg (EMBL, Heidelberg) (24) and cultured in DMEM with 10% FBS (25). All cell lines were verified by short tandem repeat (STR) profiling and tested negative for mycoplasma contamination.

### Single-molecule RNA fluorescent *in situ* hybridization (FISH)

Cells were grown on coverslips, briefly washed with PBS and fixed with PBS/3.7% formaldehyde at room temperature for 10 min. Following fixation, cells were washed twice with PBS. The cells were then permeabilised in 70% ethanol for at least 1 hour at 4°C. Stored cells were briefly rehydrated with Wash Buffer (2X SSC, 10% formamide, Biosearch) before FISH. The Stellaris FISH Probes (*lnc289* exonic probes Q570; sequences in Supplementary Table 4) were added to the hybridization buffer (2X SSC, 10% formamide, 10% dextran sulfate, Biosearch) at a final concentration of 250 nM. Hybridization was carried out in a humidified chamber at 37°C overnight. The following day, the cells were washed twice with Wash Buffer (Biosearch) at 37°C for 30 min each. The second wash contained DAPI for nuclear staining (5 ng/ml). The cells were then briefly washed with 2X SSC and then mounted in Vectashield (Vector Laboratories, H-1000). Images were captured using a Nikon TE-2000 inverted microscope with NIS-elements software, a Plan Apochromat 100x objective and an Andor Neo 5.5 sCMOS camera. We acquired 25 optical slices at 0.3 μm intervals. Images were deconvolved with Huygens Professional and projected in two dimensions using ImageJ.

### Plasmids and antibodies

Plasmids used in this study were pHR-SFFV-dCAS9-BFP-KRAB (Addgene, #46911), pU6-sgRNA EF1Alpha-puro-T2A-BFP (Addgene, #60955), second-generation packaging plasmid psPAX2 (Addgene, #12260) and the envelope plasmid pMD2.G (Addgene, #12259). pHIV-Zsgreen (Addgene #18121) and LincExpress-mCherry (modified version of pLenti6.3/TO/V5-DEST, kindly provided by John Rinn, Harvard University) were used as positive controls for transduction efficiency. Cas9 antibody was obtained from Cell Signaling (#14697, dilution 1:1000) and β-tubulin was purchased from Sigma (#T019, dilution 1:2000).

### RNAi- and LNA-mediated gene depletion

HeLa cells were transfected with Lipofectamine RNAiMax reagent (Thermo Fischer Scientific) following the manufacturer’s instructions. All experiments were done 48 hours after transfection. The siRNAs (Thermo Fischer Scientifc) and LNA Gapmers (Exiqon) were used at a final concentration of 50 nM and 25nm, respectively. siRNA and LNA sequences are listed in Supplementary Table 5 and 6, respectively.

### Western blotting

Cells were grown in a 6 well plate, trypsinized, pelleted and washed twice with PBS. The pellet was lysed in lysis buffer (50 mM Tris-HCl, pH 8, 125 mM NaCl, 1% NP-40, 2 mM EDTA, 1 mM PMSF, and protease inhibitor cocktail [Roche]) and incubated on ice for 25 min. The samples were centrifuged for 3 min at 12 000 × g and 4°C. Supernatant was collected and protein concentration was determined using the Direct Detect^®^ Spectrometer (Merck Millipore). The proteins (25 μg) were denatured, reduced, and separated with Bolt^®^ 4-12% Bis-Tris Plus Gel (Thermo Fisher Scientific) in MOPS buffer (Thermo Fisher Scientific, B0001-02). The proteins were then transferred to nitrocellulose membrane and blocked with 5% nonfat milk in TBS-T (50mM Tris, 150mM NaCl, 0.1% Tween-20) for 1 hour at room temperature. The membranes were incubated with primary antibodies in 5% milk in TBS-T. After overnight incubation at 4°C, the membranes were washed with TBS-T and incubated with HRP secondary antibodies (GE Healthcare Life Sciences, 1:5000), and immunobands were detected with a Supersignal West Dura HRP Detection Kit (Thermo-Scientific). An uncropped scan of the immunoblot (Figure 2) is shown in Supplementary figure 12.

### Time-lapse microscopy

HeLa cells (10 000 cells) were cultured in eight-well chamber slides (Ibidi) with 200 μl/well of normal HeLa medium (DMEM, 10% FBS). 30 min before live-cell imaging, the medium was replaced with imaging medium (DMEM fluorobrite, A1896701, Thermo Fisher Scientific, supplemented with 10% FBS and 4mM Glutamax) containing 300nM SiR–Hoechst (Spirochrome). SiR–Hoechst was present in the medium throughout imaging. HeLa Kyoto cells were plated in the same way but imaging was performed in DMEM medium with 10% FBS. Time-lapse microscopy was performed for both cell lines 48 hours after transfection with LNA or CRISPRi transduction. Mitotic duration was measured as the time from nuclear envelope breakdown (NEBD) until anaphase onset, based on visual inspection of the images. Live-cell imaging was performed using a Zeiss Axio Observer Z1 microscope equipped with a PL APO 0.95NA 40X dry objective (Carl Zeiss Microscopy) fitted with a LED light source (Lumencor) and an Orca Flash 4.0 camera (Hamamatsu). Four positions were placed per well and a z-stack was acquired at each position every 10 minutes for a total duration of 12 hours. Voxel size was 0.325 μm × 0.325 μm × 2.5 μm. Zen software (Zeiss) was used for data collection and analysis. Throughout the experiment, the cells were maintained in a microscope stage incubator at 37 °C in a humidified atmosphere of 5% CO_2_.

### CRISPR interference (CRISPRi) and gRNA design

For CRISPRi, we used two negative control sgRNAs and one positive control sgRNA (against the *H19* lncRNA) (8). For *lnc289*, *H19* and *ch-TOG*, we designed gRNA sequences (20 nt) targeting a genomic window of −50 to +200 bp relative to the transcription start site (TSS) (Supplementary Table 7). The location of the TSS was determined using the NCBI RefSeq database. The MIT CRISPR (http://crispr.mit.edu) and the gUIDEbook™ gRNA design (Desktop Genetics Ltd) web tools were used to design the gRNA sequence. Potential off-target effects were analysed with the MIT CRISPR and the CRISPR RGEN Cas-OFFinder web tools, for which the position of off-target alignments with equal or less than 4 mismatches were separately checked using the Basic Local Alignment Search Tool. Additional sequences were added to the sgRNA sequences to obtain compatible sticky ends for cloning the DNA insert into the 5’BstXI-BlpI3’ digested backbone of a pU6-sgRNA EF1Alpha-puro-T2A-BFP expression plasmid. gRNA oligos were phosphorylated, annealed and cloned into pU6-sgRNA EF1Alpha-puro-T2A-BFP expression plasmid. All inserts were verified with Sanger sequencing.

### Lentiviral transduction

To produce lentivirus, 4 × 10^6^ of HEK293T cells were plated in a 10 cm dish one day prior to transduction. Next day, cells were transfected with 15 μg of DNA, composed of 9 μg of the lentiviral vector DNA containing the transgene, 4 μg of psPAX.2 and 2 μg of pMD2.G in the final transfection volume of 1.5 ml (including 45 ul of Trans-Lt1 transfection reagent, Mirus) using OptiMEM medium (Thermo Fisher Scientific). As a positive control for viral infection and to control for any possible effects of lentiviral delivery, we transduced cells with polybrene (5 μg/ml, Sigma) or with pHIV-Zsgreen and LincExpress-mCherry vectors. The transfection mixture was incubated for 25 min at room temperature. Prior to transfection, old medium was replaced by 14 ml of fresh medium and transfection mix was added dropwise to the cells and incubated for 24 hours at 37°C. The following day, old medium was replaced by 7 ml of fresh medium and incubated for another 24 hours at 37°C. Viral supernatant was collected 48 and 72 hours post transfection, spun down at 1800 × g for 5 min at +4°C, and filtered through a 45 μm filter. Ready-to-use virus was stored at +4°C. For long-term storage, viral supernatant was frozen at −80°C.

### FACS analysis and cell sorting

HeLa cells were transduced with lentivirus containing the pHR-SFFV-dCAS9-BFP-KRAB vector together with polybrene (5 μg/ml, Sigma). 24 hours after lentivirus transduction, the medium was replaced and the cells were incubated for another 48 hours. HeLa cells were then sorted for the BFP-expressing cells using the BD FACSAria III cell sorter (CRUK Flow Cytometry Core Facility). The expression of BFP fluorescent proteins was detected using MACSQuant VYB (Miltenyi Biotec) and the data were analysed using the FlowJo v7.1 software. BFP-sorted HeLa cells were used for single cell cloning in 96-well plate (clonal cells) or to create a stable non-clonal cell population.

### CRISPRi-mediated depletion

Three to four days after FACS sorting, dCas9-KRAB transduced cells were plated on 12-well plates and infected with lentivirus containing gRNAs targeting *lnc289*, *ch-TOG* or *H19*, or with lentivirus containing two negative guide RNAs. The lentivirus was diluted with HeLa medium (1:1 dilution) and cationic polymer polybrene was added to facilitate viral transduction (5 μg/ml, Sigma). After a 24 hour incubation, supernatant was removed and fresh medium was added for another 48 hour incubation before RNA collection to evaluate knockdown of the target gene. Non-transduced cells did not receive virus and were used as a negative control.

### RNA extraction, cDNA and Quantitative Real-Time PCR (qPCR)

RNA (1 μg) was extracted with the RNeasy Kit (QIAGEN, 74106) and treated with DNase I following the manufacturer's instructions (QIAGEN, 79254). The QuantiTect Reverse Transcription Kit (QIAGEN, 205313) was used for cDNA synthesis including an additional step to eliminate genomic DNA contamination. Quantitative real-time PCR (qPCR) was performed on a 7900HT Fast Real-Time PCR System (Applied Biosystems) with Fast SYBR Green Master Mix (Life Technologies). Thermocycling parameters were defined as 95°C for 20 sec followed by 40 cycles of 95°C for 1 sec and 60°C for 20 sec. Two reference genes (*GAPDH* and *RPS18*) were used to normalise expression levels using the 2^-ΔΔCT^ method. Sequences of qPCR primers are provided in Supplementary Table 8.

### Subcellular fractionation

RNA was fractionated as described previously (26). Briefly, cells from a 150 mm dish were used to isolate RNA from cytoplasmic, nucleoplasmic and chromatin fractions by TRIZOL extraction (Life Technologies). Expression of target genes in each fraction was analysed by qPCR. Data were normalised to the geometric mean of *GAPDH* and *β-actin* levels in each cellular compartment. *MALAT1* and *RPS18* were used as positive controls for chromatin and cytoplasmic fractions, respectively.

### RNA library preparation, sequencing and analysis

RNA-seq libraries were prepared from HeLa cells using TruSeq Stranded Total RNA Kit with Ribo-Zero Gold (IIllumina, RS-122-2303). We performed four biological replicates for RNAi, LNAs and CRISPRi-mediated depletion of *lnc289* and *H19*. Indexed libraries were PCR-amplified and sequenced for 125bp paired-end reads on an Illumina Hiseq 2500 instrument (CRUK Genomics Core Facility). Each library was sequenced to a depth of 20-30 million read pairs. Paired-end reads were aligned to the human genome hg38 (27) and the number of read pairs mapped to the exonic regions of each gene was counted for each library (28). Approximately 80% of read pairs contained one read that was successfully mapped to the human reference genome. On average, 74% of all read pairs in each library were assigned into exonic regions and counted. Any outlier samples with very low depth (resulting from failed library preparation or sequencing) were removed prior to further analysis. Differential gene expression analyses were performed using the voom-limma framework (29), where we tested for differential expression above a log_2_-fold change threshold of 0.5 in pairwise contrasts between groups of samples. For each contrast, genes with significant differences in expression between groups were detected at a false discovery rate (FDR) of 5%.

### Determination of noncoding potential of lncRNAs

The Coding-Potential Calculator (CPC) ((30), http://cpc.cbi.pku.edu.cn) and Coding Potential Assessment Tool (CPAT) ((31), http://lilab.research.bcm.edu/cpat/index.php) were used to determine noncoding potential. LncRNAs with CPC score >1 and CPAT score >0.364 were predicted to have protein-coding capacity. The PhyloCSF score was taken from UCSC (https://github.com/mlin/PhyloCSF/wiki, (32)).

### Statistical analysis

The statistical significance of data was determined by two-tailed Student's t-test in all experiments using GraphPad Prism unless indicated otherwise. P-values > 0.05 were considered statistically not significant. The differential expression analysis is described in detail in the Supplementary Methods.

## RESULTS

### LNA and RNAi technologies are associated with non-negligible off-target effects

We first verified that RNAi (pool of four different siRNAs) and LNA oligonucleotides were able to successfully deplete both protein-coding genes and lncRNAs by knocking down *Ch-TOG/CKAP5* and *MALAT1*, respectively (Supplementary Figure 1). Both technologies routinely achieved high knockdown efficiency, indicating that gene depletion was effective.

We then performed RNA-seq on untreated cells and cells treated with one of two negative control siRNAs (Ambion and GE Dharmacon) (Figure 1A). In principle, treatment with the negative control siRNAs should have no effect on gene expression, as no gene is targeted for depletion. Thus, our experimental design allows us to quantify the transcriptional off-target effects of the two negative control siRNAs, based on the number of differentially expressed genes (DEGs) when compared to untreated cells. In this analysis, genes are only considered to be subject to off-target effects if the logfold change between conditions is significantly greater than 0.5 (33). This ensures that only modest-to-large changes in gene expression are detected, which focuses on DEGs that are more likely to have some biological effect.

**Figure 1.**
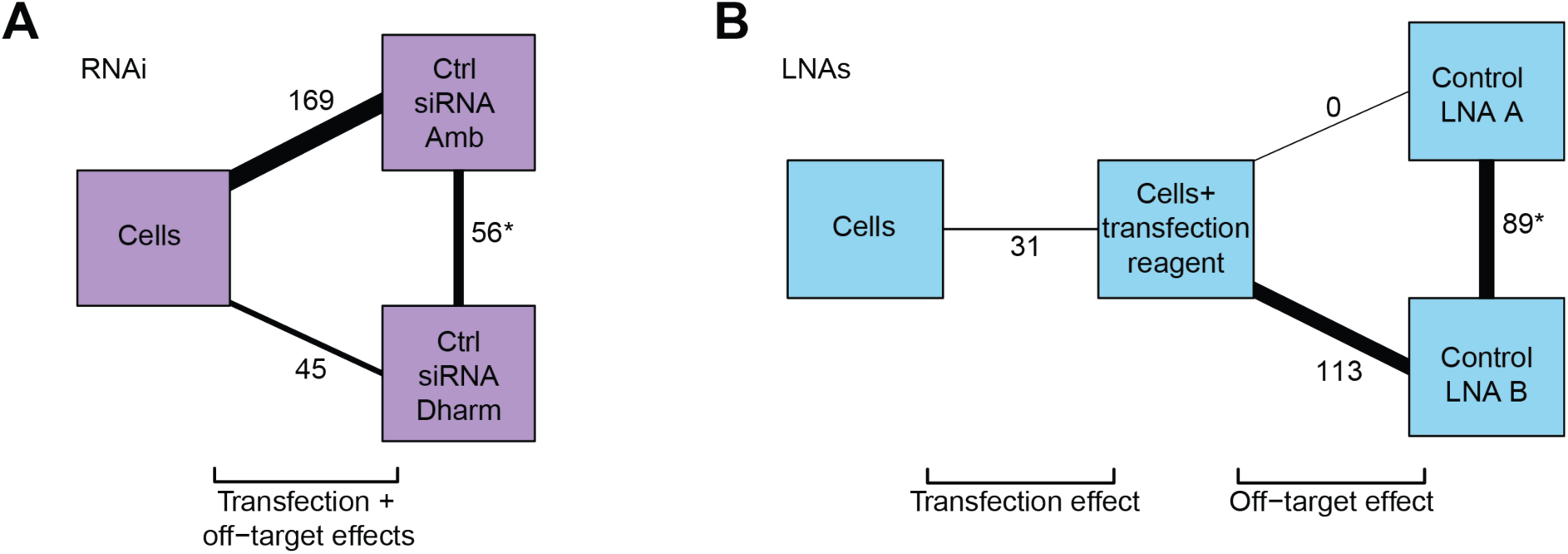
Off-target effects associated with RNAi and LNA oligonucleotides. **A.** Comparison of the transcriptional differences between untreated cells and cells treated with two negative control siRNAs (Ambion and GE Dharmacon). The number of DEGs between each pair of treatments is labelled and shown as connecting lines of proportional thickness. **B.** Comparison of the transcriptional differences between untreated cells, cells treated with transfection reagent (RNAiMax) and two negative control LNAs (A and B). The number of DEGs differing between each pair of treatments is labelled as described in **A**. DEGs for each pairwise comparison were defined at an FDR of 5% after for a log_2_-fold change significantly greater than 0.5. Lists of DEGs for each comparison are shown in Supplementary Table 1.

We identified 45 (GE Dharmacon) and 169 genes (Ambion) affected by the introduction of each negative control siRNA at a FDR of 5%, some of which we validated by qPCR (Supplementary Figure 2A). Of these, 30 genes were affected by both negative control siRNAs, likely representing a general effect of siRNA transfection (Supplementary Figure 2B). The off-target genes affected by either of the negative control siRNAs were not obviously associated with a functional pathway. No single KEGG term contained more than 20% of the DEGs (Supplementary Figure 3A and B, Supplemental Table 1), with most terms containing less than 10%. These results indicate that offtarget genes associated with addition of negative control siRNAs do not fall into pathways that would be easy to computationally predict or remove.

We further identified DEGs between two cell cultures, each of which had been treated separately with one of the two negative control siRNAs. In this comparison, only the siRNA sequence differs between the two controls – thus, any DEGs between the controls must represent sequence-dependent off-target effects, rather than a general effect of siRNA transfection. Comparison between the negative control treatments yielded 56 DEGs, indicating that the perturbations due to nonspecific targeting is dependent on the exact sequence of the siRNA used to treat the cells. Again, no common function for this set of DEGs was detectable by KEGG pathway analysis (Supplementary Figure 3C, Supplemental Table 1), with fewer than 10% of DEGs associated with any KEGG term.

We also generated transcriptomic profiles for negative controls at each step of the depletion protocol using locked nucleic acid (LNA) Gapmers (Exiqon). Treating cells only with transfection reagent led to perturbation of 31 genes (Figure 1B) compared to untreated cells. Treatment using either of two different negative control LNAs yielded zero (control LNA A, Exiqon part number 300611-00) and 113 (control LNA B, Exiqon part number 300615-00) DEGs compared to the transfection control, which we validated by qPCR (Supplementary Figure 4A). Comparison between the two negative control LNAs identified 89 DEGs. Applying the same reasoning as described above for RNAi, these 89 genes represent the typical scale of sequence-dependent off-target effects of the LNA approach. Similar to RNAi, KEGG pathway analysis of off-target effects using negative LNA controls did not reveal any common function for the DEGs (Supplementary Figure 4B and C; Supplementary Table 1).

In summary, a non-negligible number of genes display off-target activity with both RNAi and LNA technologies. The sequence-dependent nature of the off-target effects has important implications for how these methods can be used to study transcriptional regulation. In particular, the high sequence specificity of off-target effects in both methods strongly suggests that generic negative controls cannot accurately recapitulate nonspecific changes in expression that arise when targeting a particular gene.

### CRISPRi with single cell cloning introduces transcriptional variation

CRISPRi-mediated transcriptional inhibition can target gene expression using both non-clonal (8,9,15) and single cell derived clonal populations (11,15). To directly compare CRISPRi with other LOF methods, we generated HeLa cells expressing dCas9-KRAB using lentiviral transduction, and confirmed the expression of dCas9-KRAB in both clonally isolated populations of cells and non-clonally isolated populations (Figure 2A). We verified that our CRISPRi system was effective at knocking down protein-coding genes by targeting *Ch-TOG/CKAP5*, a microtubule associated protein required for mitotic spindle assembly (Supplementary Figure 5A) (34). Moreover, depletion of *Ch-TOG* with CRISPRi recapitulated known cellular phenotype such as mitotic delay (Supplementary Figure 5B) (35).

**Figure 2.**
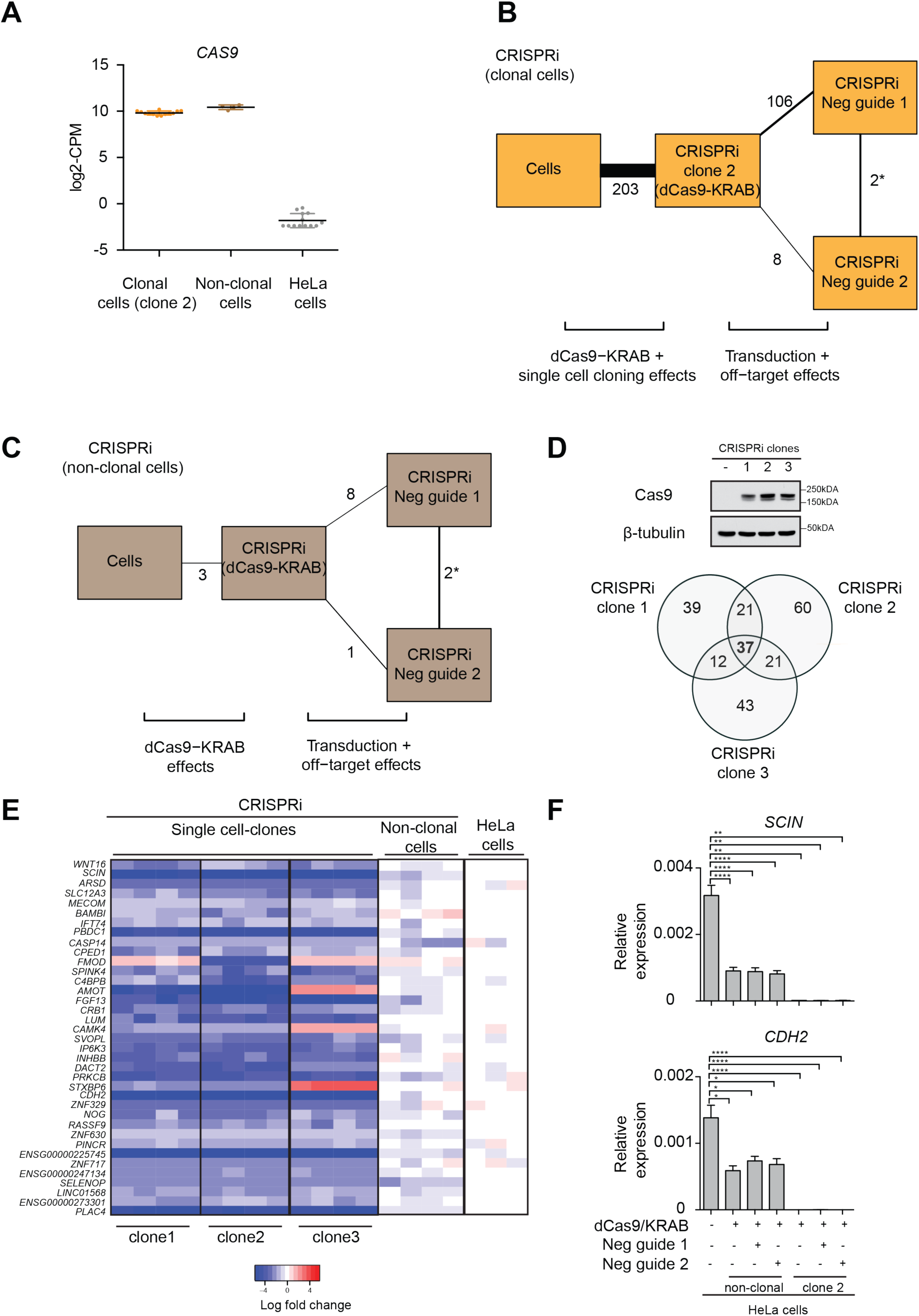
Clonal variations in CRISPRi and their associated off-target transcriptional effects. **A.** Expression in counts-per-million (CPM) of *Cas9* in CRISPRi clonal, CRISPRi non-clonal and untransduced HeLa cells. Clone 2 was used for showing *Cas9* expression in CRISPRi clonal cells. **B.** Comparison of the transcriptional differences between parental HeLa untreated cells, CRISPRi clones expressing only dCas9-KRAB (clone 2) and clones treated with two negative guide RNAs (negative guide 1 and 2). The number of genes differing between each pair of treatments is labelled and shown as connecting lines of proportional thickness. **C.** Comparison of the transcriptional differences between parental HeLa untreated cells, non-clonal CRISPRi cells expressing dCas9-KRAB and non-clonal cells treated with two negative guide RNAs (negative guide 1 and 2). The number of genes differing between each pair of treatments is labelled as described in **B**. **D.** Expression of dCas9-KRAB in three different CRISPRi clones derived from single cell cloning, confirmed by immunoblot using a Cas9 antibody. β-tubulin was used as a loading control. A Venn diagram of DEGs detected in the three different clones against untransduced cells in the absence of any guide RNAs identified 37 genes as a common transcriptional signature of cloning. The total number of genes in this analysis was 17991 and DEGs were detected at a FDR of 5%. **E.** Heat map of DEGs from three different CRISPRi clones compared to non-clonal cells and parental untransduced HeLa cells, in the absence of any guide RNAs. 33 out of 37 genes were downregulated in clonal cells compared to the parental population. **F.** Downregulation of two randomly selected DEGs from **E** (*SCIN* and *CDH2)* was validated by qPCR in clonal cells (clone 2) and in non-clonal populations. Expression levels were normalized to the geometric mean of *GAPDH* and *RPS18.* Error bars, s.e.m. (*n*=4 biological replicates). Statistical significance by two-tailed Student’s *t*-test: \**P*<0.05, \*\**P*<0.01, *** *P*<0.001 and \*\*\*\**P*<0.0001. Lists of DEGs for each pairwise comparison in **B** and **C** are provided in Supplementary Table 1.

We then transduced one CRISPRi clone (CRISPRi clone 2) and a non-clonal cell population with two negative control guide RNAs (negative control guide 1 or 2). We performed RNA-seq to quantify the off-target effects by comparing the transcriptional profile before and after treatment with each of the negative control guides. The addition of the negative guide RNAs had modest effects in the clonal CRISPRi cells (8 and 106 genes, Figure 2B) and minor effects in the non-clonal CRISPRi cells (1 and 8 genes, Figure 2C). This is noticeably lower than the extent of off-target effects associated with RNAi or LNAs, consistent with previous studies suggesting that CRISPRi-mediated gene repression is highly specific (7). Moreover, only 2 genes were differentially expressed between the negative guide RNAs in each CRISPRi strategy. These results indicate that the off-target effects of CRISPRi are minor, not specific to the guide sequence, and can be expected to cancel out when different guide RNA treatments (negative control or targeting a specific gene) are compared.

Single cell cloning after introduction of dCas9-KRAB provides another potential source of offtarget effects. Indeed, we found 203 DE genes between the parental HeLa cells and clonally-derived cells expressing dCas9-KRAB in the absence of a guide RNA (Figure 2B); in contrast, no such difference (only three genes) was observed between the parental HeLa cells and the non-clonal cells (Figure 2C). To investigate whether the single cell dilution protocol for obtaining CRISPRi targeted cell clones could be responsible for the increase in the number of DEGs, we profiled additional CRISPRi clones expressing different levels of dCas9-KRAB (clones 1 and 3; Figure 2D) along with additional independent replicates of CRISPRi clone 2. The DEG analysis yielded a core set of 37 DE genes that were consistently and strongly downregulated in all three clones compared to non-transduced control cells (Figure 2D and E). We did not observe any common pathway for these 37 genes using KEGG analysis (Supplementary Figure 6, Supplementary Table 1) and found no relationship between these genes and their chromosomal location (Supplementary Figure 7).

Most of the genes (*33* out of 37) were downregulated in all clones, suggesting that they could potentially be direct targets of the repressive KRAB domain. We validated the repression of two of these 37 genes (*SCIN and CDH2*) by qPCR, as well as in HeLa cells that were transduced with dCas9-KRAB but without single cell clonal selection (Figure 2F). Importantly, these changes in gene expression are not caused by lentiviral transduction (Supplementary Figure 8), but are a consequence of single cell cloning. Indeed, no equivalent effect was observed with non-clonal cells (Supplementary Figure 9). Thus, stable expression of dCas9-KRAB causes marked changes in the transcriptomic background in addition to the depletion of gene of interest.

Widespread genome binding and modest off-target effects have been reported for the dCas9-KRAB system (22,23,36). Therefore, we compared the published binding sites for dCas9 in HEK293T cells (37) with the gene bodies for our core set of 37 differentially expressed genes (Figure 2E), and found no overlap. We performed a similar analysis using the genome-wide mapping of dCas9-KRAB in K562 cells (23) and found only two dCas9-KRAB binding events in these 37 genes. We also found no overlap between our core set of genes with those reported as being differentially expressed upon dCas9-KRAB transduction (23). These poor overlaps may reflect the potential dependency of CRISPRi off-target effects on the cell type, the epigenetic landscape, or other factors (38). More generally, a common blacklist for *in silico* removal of likely affected genes is unlikely to be effective. The absence of a consistent set of genes across three independent studies that have attempted to identify common CRISPRi off-targets suggests that purely computational approaches may not be able to account for off-target effects *a priori*.

### A case study in using LOF methods to deplete a nuclear long noncoding RNA

We applied the three most commonly used LOF methods – RNAi, LNA oligonucleotides and CRISPRi – to study the regulatory function of a previously uncharacterised lncRNA in HeLa cells. This represents a common use of LOF strategies, given that tens of thousands of lncRNAs exist in the mammalian genome (39,40). Many of these are involved in regulation of diverse cellular processes acting at the transcriptional, post-transcriptional and post-translational level (41-43), but the function of the majority of lncRNAs is unknown. LNAs have been shown to be particularly efficient for depletion of nuclear lncRNAs (44-46). The CRISPRi system has also been used successfully to deplete lncRNAs in a large-scale screen (12). In specific cases, RNAi was also shown to be effective in depleting lncRNAs (26,47), including nuclear lncRNAs (most likely due to the presence of active RNAi machinery in the nucleus (48)).

We selected a prototypical lncRNA following published guidelines (10) with specific characteristics, including: (i) previously uncharacterized, (ii) consistently expressed at more than one molecule per cell, (iii) low coding potential, (iv) chromatin hallmarks of active transcription, and (v) nuclear localization. Using these criteria, we chose *loc100289019* (also known as *SLC25A25-AS1*) as a spliced and functionally uncharacterized intragenic lncRNA with three promoters and a single 3’ polyadenylation site (Figure 3A); hereafter called *lnc289.* Among all ENCODE cell lines, *lnc289* was most highly expressed in the nucleus of HeLa cells (Figure 3B). We confirmed that *lnc289* has low protein coding potential using PhyloCSF, Coding-Potential Calculator (CPC) and Coding-Potential Assessment Tool (CPAT) (Figure 3C). Furthermore, computational analysis of previously published ribosomal occupancy data indicated no translation of the *lnc289* transcript in HeLa cells (49) (Figure 3A, Riboseq track). Nevertheless, the genomic locus was actively transcribed based on the presence of both histone H3 lysine 4 trimethylated (H3K4me3) and histone H3 lysine 27 acetylated (H3K27ac) histones at the predicted *lnc289* locus (Figure 3A). We experimentally confirmed nuclear localization of *lnc289* by single-molecule RNA FISH (Figure 3D), and demonstrated by cellular fractionation that *lnc289* is enriched in chromatin (Figure 3E). We note that the vast majority of the lncRNAs currently under active investigation satisfy our selection criteria above. This suggests that our experimental evaluation of the three most widely used LOF methods is likely to be relevant to most studies of lncRNA function.

**Figure 3.**
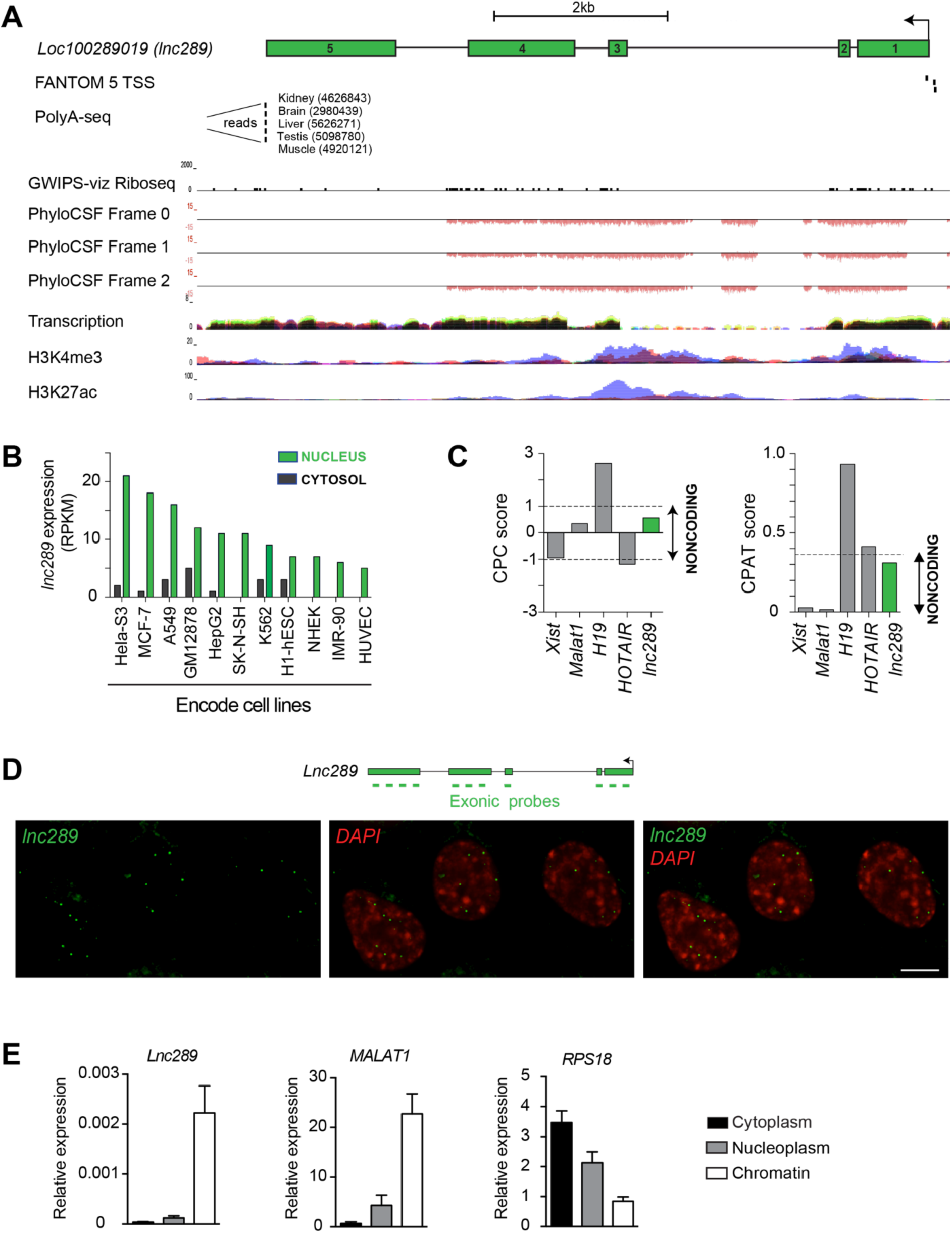
*lnc289* is an archetypical lncRNA expressed in the nucleus. **A.** Schematic representation of the genomic landscape surrounding *lnc289* (annotated in RefSeq as *loc100289019* or *SLC25A25-AS1;* chr9:128108581-128118693, hg38), including three transcriptional start sites (80) and a polyadenylation site (81). *Lnc289* is not occupied by ribosomes (49), shows no protein coding potential (PhyloCSF, (32)), and has clear hallmarks of active transcription in HeLa cells (H3K4me3 and H3K27ac data sets obtained from ENCODE via the UCSC browser). The arrows denote the direction of transcription, and green boxes represent the five exons. Note that all PhyloCSF scores at this locus are negative. **B.** Expression of *lnc289* in cytosol and nuclei of ENCODE cell lines (www.ebi.ac.uk/gxa/home), shown as reads per kilobase of exon per million reads mapped (RPKM). **C.** Computational analysis of the mature *lnc289* transcript using the CPC and CPAT tools reveals *lnc289* has low coding potential. **D.** Nuclear localization of *lnc289* in HeLa cells was determined using single-molecule RNA FISH with exonic probes (green). The nucleus was stained with 4,6-diamidino-2-phenylindole (DAPI). Scale bar represents 5 μm. **E.** *lnc289* is enriched in chromatin of HeLa cells. RNA distribution in the cytoplasm, nucleoplasm and chromatin was quantified by qPCR, and *RPS18* and *MALAT1* were used as positive controls for the cytoplasmic and chromatin fraction, respectively. Error bars represent the standard error of the mean (s.e.m) values of four independent experiments.

We then used the different LOF methods to identify the transcript(s) robustly regulated by *lnc289*, regardless of the method used for depletion. Only modest reduction of *lnc289* levels was observed with RNAi, while the other methods were able to successfully reduce *lnc289* levels by at least 50% (Figure 4 A and B). As such, we discarded the RNAi results and attempted to identify a common set of DEGs that were detected with LNA and CRISPRi methods (clonal and non-clonal cells). Remarkably, the only transcript common to these methods was *lnc289* itself (Figure 4C, Supplemental Table 2). However, the depletion of *lnc289* with LNA resulted in hundreds of DEGs, whereas many fewer DEGs were observed with CRISPRi-based methods.

**Figure 4.**
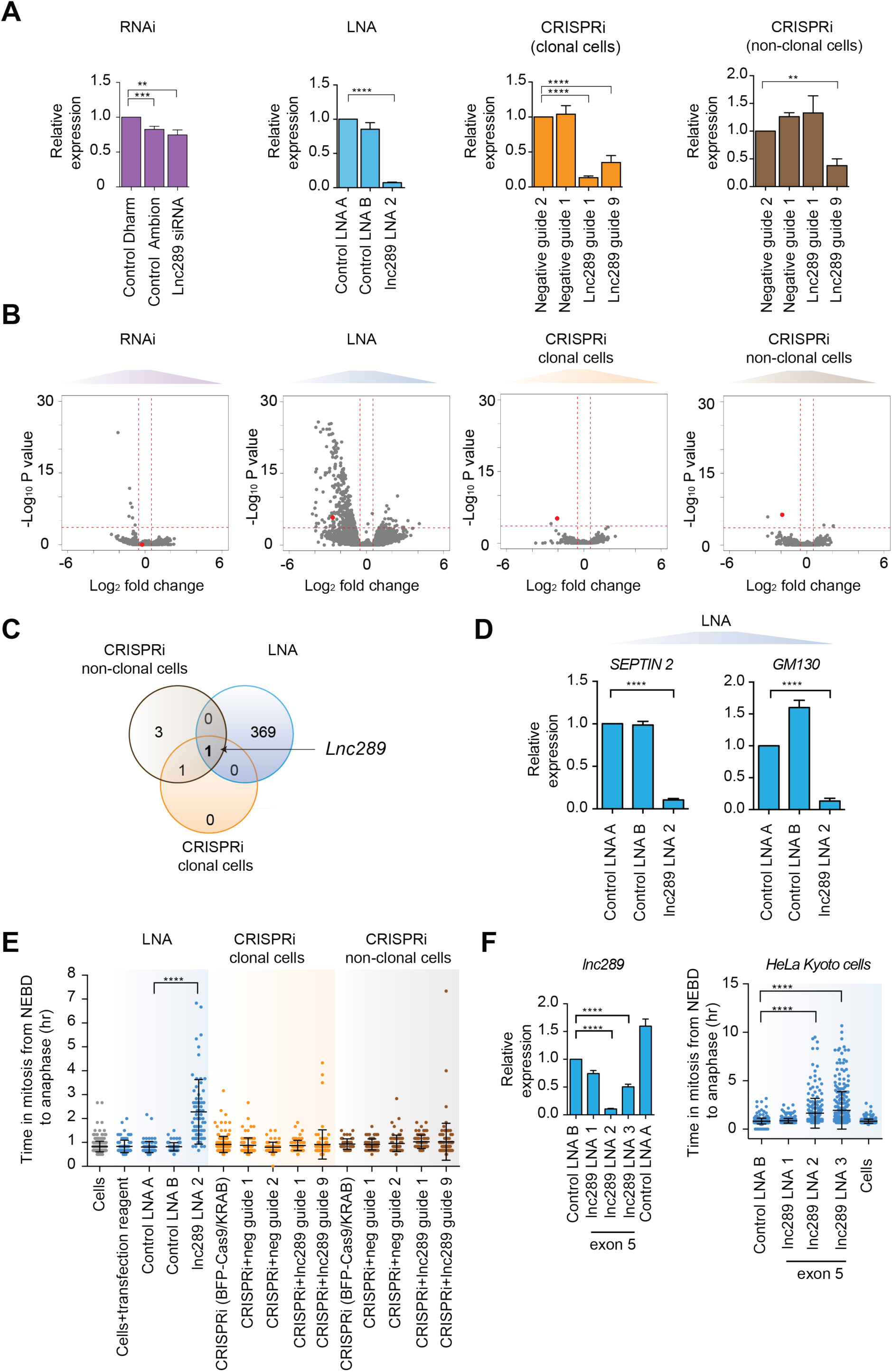
No overlap in DEGs between the different LOF methods upon depletion of nuclear *lnc289*. **A.** Expression levels of *lnc289* after RNAi, LNA and CRISPRi-mediated depletion. qPCR revealed only a 25% reduction in *lnc289* transcription after siRNA-mediated knockdown relative to negative control siRNA from Dharmacon (Control Dharm), and no significant difference relative to negative control siRNA from Ambion (Control Ambion). LNA-mediated knockdown of *lnc289* was performed using LNA oligonucleotide sequence 2 (LNA 2), and showed 90% reduction. CRISPRi-mediated repression of *lnc289* using two guide RNAs targeting TSS of *lnc289* relative to the negative (nontargeting) guide RNA 2 yielded 70-90% knockdown in clonal cells. Only one guide RNA (guide 9) was efficient in depleting *lnc289* in non-clonal cells. Statistical significance by two-tailed Student’s *t*-test: ** P<0.01, *** P<0.001 and **** P<0.0001. For all graphs, expression levels of *lnc289* were measured by qPCR using primers spanning exons 1-4, and normalized to the geometric mean of *GAPDH* and *RPS18.* Error bars, s.e.m. (*n*=4 biological replicates). **B.** Volcano plots of transcriptional differences induced by RNAi, LNA and CRISPRi-mediated depletion of *lnc289.* After siRNA-mediated depletion of *lnc289*, seven genes were differentially expressed compared to the negative control siRNAs. LNA-mediated depletion with LNA 2 identified 370 DEGs compared to negative control oligonucleotides. CRISPRi-mediated depletion of *lnc289* using guide RNA 1 and 9 revealed only two DEGs compared to negative guide RNA 2. In non-clonal cells, only four DEGs were identified using guide RNA 9 compared to the negative guides. The red horizontal line represents the significance threshold corresponding to an FDR of 5%. Red vertical lines are log_2_-fold change thresholds of +/- 0.5. The red dot corresponds to the *lnc289* itself. **C.** Venn diagram showing no overlap between the sets of DEGs identified after using LNA and CRISPRi to deplete *lnc289.* The only gene in common between LNA and CRISPRi-mediated depletion is *lnc289* itself. The total number of genes used for this analysis was 17837. **D.** qPCR confirmation of the downregulation of two DEGs (*SEPTIN2* and *GM130*) identified in **B** after LNA-mediated depletion of *lnc289* with LNA 2. Expression levels were normalized to the geometric mean of *GAPDH* and *RPS18*. Error bars, s.e.m. (n=3 biological replicates). Statistical significance by two-tailed Student’s *t*-test: \*\*\*\**P*<0.0001. **E.** Quantification results from time-lapse microscopy of mitotic progression of HeLa cells incubated with Sir-Hoechst after LNA and CRISPRi-mediated (clonal and non-clonal) depletion of *lnc289.* Mitotic duration was measured from nuclear envelope breakdown (NEBD) to anaphase onset. Bars show mean±s.d. (*n*=2 independent biological replicates). Statistical significance by Mann-Whitney test: \*\*\*\**P*<0.0001. **F.** *Left panel:* Expression analysis of *lnc289* by qPCR using two additional LNAs targeting *lnc289* (LNA 1 and LNA 3). Expression levels were normalized to the geometric mean of *GAPDH* and *RPS18.* Error bars, s.e.m. (*n*=3 biological replicates). Statistical significance by two-tailed Student’s *t*-test: \*\*\*\**P*<0.0001. *Right panel:* Quantification results from time-lapse microscopy using HeLa Kyoto cells after depletion of *lnc289* using three different LNAs. Bars show mean±s.d. (n=2 biological replicates). Statistical significance by Mann-Whitney test: \*\*\*\**P*<0.0001. The list of DEGs identified after RNAi, LNA and CRISPRi-mediated depletion of *lnc289* are shown in Supplementary Table 2.

The fact that *lnc289* is the only gene transcriptionally impacted by depletion with LNA and CRISPRi suggests two possible explanations for our results: i) *lnc289* has no function in transcriptional regulation, the lack of detection with the CRISPRi-based methods is correct, and the DE genes identified with LNA knockdown are off-target effects; or ii) *lnc289* does regulate the transcription of other genes, the LNA method correctly identifies its downstream targets, and the CRISPRi-based methods are somehow failing to recapitulate the effect.

These two explanations are mutually exclusive and cannot be easily distinguished, as the underlying problem stems from the deficiencies of our experimental tools for perturbing the biological system. We could, perhaps, infer that option (i) might be more likely if the number of DE genes after LNA depletion of *lnc289* were comparable to the number of off-target genes. However, the former (370, Figure 4B) is four times larger than the latter (89, Figure 1B), which would require large variation in the extent of the off-target effects between oligonucleotide sequences. Option (ii) is equally unappealing as it requires complex mechanisms e.g., compensatory effects in CRISPRi-depleted cells to mask the cellular phenotype (1) (50), or differences in the aspects of lncRNA biology that are disrupted by LNA and CRISPRi (1) (51).

We also examined whether the changes to the transcriptome might impact the *lnc289* knockdown phenotype. We noticed that two of the top DEGs upon LNA-mediated knockdown of *lnc289*, namely *SEPTIN2* and *GM130* (Figure 4D), had known roles in mitosis (52,53). Depletion of *SEPTIN2* leads to mitotic delay and incomplete cytokinesis whereas inhibition of *GM130* function by antibody injection leads to mitotic delay and multipolar division. Therefore, we assayed if mitotic delay also occurred as a result of the off-target effects upon *lnc289* depletion. To investigate this, we quantified the time required for HeLa cells to transition through mitosis before and after knockdown of *lnc289.* We observed a significant mitotic delay after LNA-mediated knockdown of *lnc289* (Figure 4E), whereas no such effect was observed with the CRISPRi-based methods. We further confirmed the mitotic delay with additional LNA oligonucleotide (Figure 4F) in HeLa Kyoto cells, a cell line stably expressing EGFP-alpha tubulin and mCherry-histone H2B (24). This demonstrates that the differences in the genes disrupted by each method have real consequences on the inferred biological function. Using LNA oligonuclotides, one might conclude that *lnc289* regulates mitosis, whereas the same conclusion cannot be made with CRISPRi.

We further tested CRISPRi to deplete *H19*, a multifunctional and well-characterized lncRNA with activity in the nucleus and in the cytoplasm (54)(55). Knockdown efficiency was similar in clonal and non-clonal cells (Supplementary Figure 10A), though there were modest differences in the number of DEGs by RNA-seq. Specifically, we observed 5 and 29 DEGs in clonal and non-clonal populations, respectively, compared to cells treated with negative guides (Supplementary Figure 10B; Supplementary Table 3). This difference hints at the presence of compensatory mechanisms in clonal cells that may be countering dCas9-KRAB activity, possibly as a result of the altered transcriptional background (Supplementary Figure 9B). We also examined the expression of genes previously reported to be regulated by *H19* from experiments using RNAi in different human cell lines (56) (57) (58) or genetic deletion in mice (59), but we observed no evidence of differential expression for these genes in HeLa cells after *H19* depletion (Supplementary Figure 10C). This highlights one challenge in using CRISPRi-mediated depletion to infer lncRNA function, as they often operate in a cell type-specific manner (12) (45).

## DISCUSSION

Here, we systematically compared three widely used LOF methods and evaluated the transcriptome-wide changes attributable to each individual method. We describe off-target effects associated with each LOF method that need to be considered when investigating gene function, consistent with previous studies (18-20,22,23,36,60). In particular, we identified large off-target effects in the RNAi and LNA methods, which were highly dependent on the siRNA or LNA oligonucleotide sequence. While CRISPRi was less sensitive to the guide sequence, the introduction of dCas9-KRAB provides another source of off-target effects that can significantly change the transcriptional context in the depleted cells. Single cell cloning of dCas9-KRAB-expressing cells results in strong transcriptional changes even in the absence of guide RNAs, indicating that polyclonal populations should be used for CRISPRi experiments.

Differences between the three LOF methods can also lead to significant differences in the molecular or cellular phenotype after depletion of a gene of interest, as observed in our case study with a nuclear lncRNA *lnc289*. Our results are consistent with previous studies of lncRNA depletion in mammalian cells (61,62) and zebrafish (63,64). Different strategies to deplete lncRNAs in mice have also yielded different phenotypes (65,66). Such discrepancies are not limited to lncRNAs, but have been observed when depleting protein-coding genes using RNAi and CRISPR-based methods (8,51,67,68) as well as in high-throughput screens (51) (69).

Our results suggest that CRISPRI in non-clonal populations of dCas9-KRAB-expressing cells provides the cleanest depletion of the target gene, with the fewest off-target effects (sequence-dependent or otherwise). This is consistent with previous studies demonstrating the superiority of CRISPRi compared to RNAi (7,12,60,69). However, CRISPRi has a number of limitations, especially when investigating the function of lncRNAs. Currently, CRISPRi can not differentiate *cis*- and transacting functions of RNA transcripts (41), cis-mediated regulation related to lncRNA transcription (70-72) and/or enhancer-like functions of some lncRNA loci (73-75). In addition, CRISPRi cannot be used to target bidirectional promoters (76). Similarly, CRISPRi is not ideal for targeting lncRNAs near other transcriptional units (14), as neighboring genes may be unintentionally repressed.

For studies that use LOF methods to characterize the regulatory roles of targeted genes, our recommendations are to generate libraries (1) from controls obtained at each step of the method and (2) from multiple negative control sequences. This allows accurate quantification of the extent of transcriptional off-target effects introduced upon sequential manipulations in the LOF protocol. Affected genes can then be excluded from the DE analysis in samples where the gene of interest has been depleted, thus reducing the impact of off-target effects on the biological conclusions. It may also be necessary to discard experiments where the number or log-fold changes of the off-target genes are comparable to or greater than the number or log-fold changes of DEGs detected upon knockdown of the target gene. In such cases, there is no meaningful way to distinguish between off-target effects and genuine knockdown effects.

Furthermore, we recommend performing differential expression analyses with a minimum logfold change threshold, in order to avoid detecting genes with small changes in expression. Indeed, when we repeated the analyses using a test for any differential gene expression (i.e., without a minimum log_2_-fold change threshold), we obtained a far greater number of affected genes in all comparisons for all depletion methods (Supplementary Figure 11). In particular, we observed a large increase in the number of genes detected in comparisons between control groups for all methods, corresponding to a disproportionate increase in the scale of the off-target effects. The use of a log-fold change threshold mitigates the effect of non-specific activity on the results for each LOF method, by focusing on larger and arguably more biologically relevant effects of depletion.

Finally, the prevalence of off-target effects in most of the LOF methods emphasizes the need for rigorous validation of putative downstream targets of the target gene, such as independent rescue experiments. In the case of RNAi, off-target effects could be reduced by using alternative approaches such as siPOOL (77) or C911 mismatch siRNAs (78). It remains to be seen to what extent other CRISPR genome editing strategies, including whole locus or promoter deletion, insertion of transcriptional termination sites into the gene body as well as CRISPR-Cas13 system (79), may have off-target effects that impact functional characterization of gene of interest.

Modern LOF technologies allow researchers to deplete any transcript of interest in a variety of biological systems, and provide an essential experimental toolkit for studying the biological function of transcripts and dissecting networks of transcriptional regulation. Here, we have empirically formulated recommendations to minimize technical artefacts and avoid - or at least prudently manage - the offtarget effects of these commonly used LOF methods.

## DATA AVAILABILITY

Sequencing data are available in the ArrayExpress database (http://www.ebi.ac.uk/arrayexpress) under the accession number E-MTAB-5308.

## FUNDING

This work was made possible by funding from Cancer Research UK (C14303/A17197 to F. G and A20412 to D.T.O). We also acknowledge the support of the University of Cambridge, the Wellcome Trust Investigator Award (D.T.O), European Research Council (615584; D.T.O) and Hutchison Whampoa Limited.

### Authors’ contributions

Conception and design of study: LS, ATLL, FG and DTO. Data acquisition: LS, ATLL, JM, PM, ARB, VQ. Data analyses and interpretation: LS, ATLL, FG, JCM, CB, DTO. Writing the paper: LS, ATLL and DTO with input from all of the authors wrote the manuscript.

## ACKNOWLEDGEMENTS

We thank all the members of Odom, Gergely and Marioni groups for helpful discussions. We also thank the Genomics, FACS and Microscopy Core Facilities at the CRUK Cambridge Institute and John Rinn for providing us with lincExpress vector.

## SUPPLEMENTARY DATA

Supplementary Data are available at NAR online.

